# Caspase-dependent cleavage of DDX21 suppresses host innate immunity

**DOI:** 10.1101/2021.04.06.438751

**Authors:** Wei Wu, Yang Qu, Shengqing Yu, Sa Wang, Yuncong Yin, Qinfang Liu, Chunchun Meng, Ying Liao, Zaib Ur Rehman, Lei Tan, Cuiping Song, Xusheng Qiu, Weiwei Liu, Chan Ding, Yingjie Sun

## Abstract

DEAD (Glu-Asp-Ala-Glu)-box RNA helicases have been proven to contribute to antiviral innate immunity. DDX21 RNA helicase was identified as a nuclear protein involved in ribosomal RNA processing and RNA unwinding. DDX21 was also proved to be the scaffold protein in the complex of DDX1-DDX21-DHX36 which senses double strand RNA and initiates downstream innate immunity. Here, we identified that DDX21 undergoes caspase-dependent cleavage after virus infection and treatment with RNA/DNA ligands, especially for RNA virus and ligands. Caspase-3/6 cleave DDX21 at D^126^ and promotes its translocation from the nucleus to the cytoplasm in response to virus infection. The cytoplasmic cleaved DDX21 negatively regulates the IFN-β signaling pathway by suppressing the formation of DDX1-DDX21-DHX36 complex. Thus, our data identify DDX21 as a regulator of immune balance and most importantly uncover a potential role of DDX21 cleavage in the innate immunity response towards virus.

**Importance:** Innate immunity serves as the first barrier against virus infection. DEAD (Glu-Asp-Ala-Glu)-box RNA helicases, originally considered to be involved RNA processing and RNA unwinding, have been shown to play an important role in anti-viral innate immunity. The precise regulation of innate immunity is critical for the host because the aberrant production of cytokines leads to unexpected pathological consequences. Here, we identified DDX21 was cleaved at D^126^ by virus infection and treatment with RNA/DNA ligands via the caspase-3/6-dependent pathway. The cytoplasmic cleaved DDX21 negatively regulates the IFN-β signaling pathway by suppressing the formation of DDX1-DDX21-DHX36 complex. In sum, our data identify DDX21 as a regulator of immune balance and most importantly uncover a potential role of DDX21 cleavage in the innate immunity response towards virus.

## Introduction

The host innate immunity response is initiated by virus infection. Several pathogen recognition receptors (PRRs) were mobilized to sense viral nucleic acids and ultimately lead to the induction of interferons (IFNs) and other inflammatory cytokines to protect host cells (1). Among them, endosomal toll-like receptor 3 (TLR3) (2, 3), cytoplasmic retinoic acid-inducible gene I (RIG-I), and melanoma differentiation-associated protein 5 (MDA5) (4, 5) were demonstrated to be critical for sensing viral double-stranded RNA (dsRNA). The adaptor protein TIR domain-containing adaptor inducing interferon-β (TRIF) (2) and mitochondrial protein MAVS (VISA, IPS-1, or Cardif) (6–9) then activated, leading to the activation of nuclear factor kappa B (NF-κB) and transfection factor interferon regulatory factor 3 (IRF3) as well as the production of various cytokines, including type I-IFNs (10). The secreted IFN binds to the IFN receptor and induces the expression of various interferon-stimulated genes (ISGs) to establish a cellular anti-viral state (1).

The helicase family is a class of enzymes that are essential to all living organisms. Their main function is separating nucleic acid strands (DNA, RNA, or RNA:DNA hybrid) (11). The human genome encodes 64 RNA helicases and 31 DNA helicases, which are classified in several superfamilies (SFs) based on their conserved motifs (12, 13). DEAD/H box helicase belongs to the SF2 family, the largest group of helicases, which are involved in various cellular processes (12, 13). RNA helicases are critical for most RNA metabolism processes and are also involved in the antiviral immune response by sensing foreign RNAs (14). RIG-I, MDA5, and laboratory of genetics and physiology 2 (LGP2), three RIG-I-like receptors (RLRs) that belong to SF2 RNA helicases, are closely related to DEAD-box helicases (15). In addition to RLRs, a growing list of DEAD/H box helicases has been identified to contribute to antiviral innate immunity in recent years, either by acting as sensors for viral nucleic acids or as mediators of downstream signaling events (16–21).

DDX21 was shown to be an abundant nuclear protein in HeLa cells that directly binds rRNAs and snoRNAs and promotes rRNA transcription, processing, and modification (22–25). Another important function of DDX21 is unwinding RNAs, including dsRNA and RNA guanine quadruplexes (26–28). Recently, several reports indicated that DDX21 also plays a role in innate immunity and virus infection. DDX21, together with DDX1 and DHX36, can bind the adaptor protein TRIF to sense dsRNA (20). During dengue virus infection, DDX21 translocates from nucleus to cytoplasm and mediates innate immunity (29). Moreover, DDX21 inhibits influenza A virus replication but is counteracted by viral NS1 protein (30). However, the way in which DDX21 precisely regulates anti-viral innate immunity and whether DDX21 undergoes protein modification during virus infection remain unclear.

Our preliminary screening results showed that Newcastle disease virus (NDV) manipulates the expression of several DDX/DHXs during infection (data not shown). Among them, DDX21 was found to be cleaved in HeLa cells infected with NDV. Here, we report that virus infection and RNA/DNA ligands cleave DDX21 at D^126^ via the caspase-3/6 pathway. The cleavage of DDX21 promotes its translocation from the nucleus to the cytoplasm in response to virus infection. The cytoplasmic cleaved DDX21 negatively regulates the IFN-β signaling pathway by suppressing the formation of DDX1-DDX21-DHX36 complex. Our study therefore reveals a role of DDX21 in the regulation of anti-viral innate immunity and provides molecular insights into how the host balances anti-viral and aberrant innate immunity.

## Materials and Methods

### Reagents and antibodies

Caspase inhibitor z-VAD-FMK (C1202, Beyotime, Nantong, China) was used at a concentration of 50 μM. Neddylation inhibitor MLN4924 (S7109, Selleckchem, Houston, TX, USA) was used at a concentration of 1 μM. Proteasome inhibitor MG-132 (S1748, Beyotime) was used at a concentration of 20 μM. The autophagy inhibitors wortmannin (W1628, Sigma-Aldrich, St. Louis, MO, USA) and chloroquine (CQ) (C6628, Sigma-Aldrich) were used at 300 nM and 25 μM concentrations, respectively. Poly(I:C) (tlrl-pic), 3p-hpRNA (tlrl-hprna), HSV-60 (tlrl-hsv60n) and poly(dG:dC) (tlrl-pgcn) were purchased from Invivogen (San Diego, CA, USA). Rabbit monoclonal anti-DDX21 (ab182156), anti-caspase-6 (ab185645), mouse monoclonal anti-VSV-glycoprotein (G) antibodies (ab50549), and rabbit polyclonal anti-Lamin B1 antibodies (ab16048) were purchased from Abcam (Cambridge, MA, USA). Rabbit polyclonal anti-HSV-1-glycoprotein D (gD) (NB600-516) was purchased from Novus biologicals (Littleton, CO). Rabbit polyclonal anti-Sendai (PD029C1) was purchased from MBL (Nagoya, Japan). Rabbit polyclonal anti-caspase-3 (GTX110543) was purchased from GeneTex (Irvine, CA). Mouse monoclonal anti-Flag (F1804), anti-HA (H9658), and anti-β-actin antibody (A1978) were purchased from Sigma-Aldrich. Rabbit monoclonal anti-phospho-TANK Binding Kinase 1 (TBK1) (#5483) and anti-TBK1 (#3013) were purchased from Cell Signaling Technology (Beverly, MA, USA). Monoclonal antibody against NDV nucleoprotein (NP) was prepared in our laboratory (31). For the immunofluorescence assays, mouse monoclonal anti-HSV-1/2 gE was purchased from Santa Cruz Biotechnology (Dallas, TX, USA), and mouse polyclonal anti-VSV-G antibodies (ab1874) were purchased from Abcam. IFN-β was measured with an ELISA kit (41410, PBL Assay Science, Piscataway, NJ).

### Cell cultures and virus

HeLa, A549, HEK-293T, Vero, HuH7, and THP-1 cells were purchased from American Type Culture Collection (ATCC). These cells were maintained in Dulbecco’s modified Eagle’s medium (DMEM) supplemented with 10% fetal bovine serum (Thermo Fisher Scientific, Waltham, MA, USA). NDV Herts/33 strain was obtained from the China Institute of Veterinary Drug Control (Beijing, China). Herpes simplex virus (HSV)-1 was kindly provided by Yasushi Kawaguchi (University of Tokyo, Japan), and vesicular stomatitis virus (VSV) was provided by Jianchao Wei (Shanghai Veterinary Research Institute, China). Sendai virus (SeV) was provided by Quan Zhang (Yangzhou University, China). HSV-1 and VSV titration was determined as the median tissue culture infective dose (TCID_50_) on Vero cells.

### Plasmids

Flag-tagged DDX21-X1 (Flag-DDX21) and DDX21-X2 were constructed by inserting the open reading frame (ORF) of human DDX21 isoform 1 (NM_004728.4) and isoform 2 (NM_001256910.1) into plasmid p3XFLAG-CMV-14 (Sigma-Aldrich), respectively. Flag-tagged deletion constructs (Δ217-395, Δ396-573) and point mutants of DDX21 (D87A, D126A, 160A, D87A/D126A, D87A/160A, D126A/D160A, D81A/D126A/160A) were generated by site-directed mutagenesis, as described previously (32, 33). Flag- and HA-tagged wide-type and truncates of DDX21 (1-125, 127-784, Δ1-216, Δ574-784) were constructed by inserting indicated sequences into p3XFLAG-CMV-14 (Sigma-Aldrich) and pCMV-HA (Promega), respectively. HA-tagged DDX1and HA-tagged TRIF were constructed by inserting indicated sequences into pCMV-HA (Promega), respectively. Myc-tagged DHX36 was constructed by inserting the ORF of DHX36 into pCMV-Myc (Promega). pHAGE-WT and -D126A DDX21 were constructed by inserting the flag-tagged WT and D126A DDX21 into pHAGE-bsd, which was constructed based on pHAGE-puro (Addgene, plasmid 118692). The primer sequences for plasmid construction are listed in Supplementary Table 1. The IFN-β promoter luciferase reporter was kindly provided by Takeshi Fujita (Kyoto University, Japan).

### Cell transfection, luciferase assay, and gene knockdown

Cells were transfected using FuGENE® HD (Promega, Madison, WI, USA) or Lipofectamine® 2000 (Thermo Fisher Scientific) according to the manufacturer’s instructions. For the luciferase assay, cells were cultured in 24-well plates and co-transfected with 100 ng of firefly luciferase reporter (IFN-β-Luc) and 10 ng of the constitutive *Renilla luciferase* reporter pRL-TK. Luciferase activity was measured at 24 h post transfection (hpt). Lentiviral shRNA for targeting endogenous DDX21 (5’-CCCATATCTGAAGAAACTATT-5’) were purchased from Gene Pharma (Shanghai, China). To generate DDX21 stable knockdown cell line, HeLa cells were infected with lentiviral shDDX21 and selected by puromycin as described previously (34).

### Immunofluorescence (IF) assay

HeLa cells were washed in PBS, fixed in 4% neutral formaldehyde, and then permeabilized with 0.5% Triton X-100 in Tris-buffered saline with Tween 20 (TBST) for 10 min. After blocking in TBST with 3% bovine serum albumin, the cells were incubated with primary antibody for 1 h at 37°C. The cells were washed three times with TBST and incubated with secondary antibody. Subsequently, they were washed and incubated with another primary antibody, followed by incubation with secondary antibody. Next, the cells were washed again and incubated with 0.5 μg/mL DAPI. The coverslips were washed and visualized using an LSM 880 Zeiss confocal microscope (Carl Zeiss, Jena, Germany).

### Nucleocytoplasmic separation assay

Nuclear extractions were prepared using NE-PER nuclear and cytoplasmic extraction reagents (78833, ThermoFisher Scientific) according to the manufacturer’s instructions.

### Immunoblotting and co-immunoprecipitation

Immunoblotting was performed as described previously (31). Briefly, cells were lysed in cell lysis buffer containing a protease inhibitor cocktail (Merck Millipore, Darmstadt, Germany). The lysates were denatured and then subjected to sodium dodecyl sulfate polyacrylamide gel electrophoresis (SDS–PAGE) and immunoblotting and quantified using Image J software. For co-immunoprecipitation, HeLa cells were transfected with expression vectors for 24 or 36 h and lysed with a cell lysis buffer (150 mM NaCl, 50 mM Tris-HCl, pH 8.0, 5 mM EDTA, 0.5% NP-40) containing 1 mM PMSF and protease inhibitors (Merck-Millipore). The lysates were centrifuged at 12,000 g for 10 min and precipitated with anti-Flag antibody, in conjunction with Protein G Agarose beads (ThermoFisher Scientific) overnight at 4°C. The beads were washed with lysis buffer four times and eluted with SDS loading buffer by boiling for 10 min and then subjected to immunoblotting.

### Quantitative real-time PCR

Quantitative real-time PCR (qRT-PCR) was performed as described previously (35). Briefly, total RNA was extracted, reverse transcribed to cDNA, and subjected to qRT-PCR analysis using Premix Ex Taq reagents (Takara, Dalian, China). Primers were referenced from previous reports (35) (Supplementary Table 1). The relative abundance of mRNAs was calculated using the comparative C_T_ (ΔΔC_T_) method (36). All experiments were carried out in triplicate.

### Generation of knockout cells

The CRISPR/Cas9 system was used to generate *ddx21* and *casp6* knockout cells. The guide RNA (gRNA) specific to *ddx21* and *casp6* gene was designed using the online CRISPR design tool (http://crispr.mit.edu/). The gRNA oligonucleotides were annealed and cloned into the pGK1.1 (Geneloci, Nanjing, China). HeLa cells were electrotransfected with the plasmid at 550V, 1 pulse. After 24 h of electrotransfection, the supernatants were replaced with 10% FBS DMEM supplemented with 1 μg/ml puromycin (Merck-Millipore) for 24 h. The pool (mixed clones) cells were preliminarily sequenced and validated. The candidate positive clones were then subcloned onto 96-well plates using the limiting dilution method. The genomic region surrounding the CRISPR target site was amplified by PCR using the check primers. The primer sequences for sgRNAs and check primers are listed in Supplementary Table 1. The clones were sequenced to ensure the frameshifting mutation of both alleles of the established cell line. Except for PCR verification, the cells were re-checked by immunoblotting using DDX21 antibody. *Casp3-/-* HeLa cells were purchased from EdiGene Inc. (Beijing, China) and verified by immunoblotting. Casp3/6 double knockout cells were constructed by transfecting *casp6* sgRNA into *casp3-/-* HeLa cells as described above.

### Generation of cells stably expressing WT and mutant DDX21

HEK-293T cells were transfected with pHAGE-WT and -D126A DDX21, together with two package plasmids, pMD2.G (Addgene, plasmid 12259) and psPAX2 (Addgene, plasmid 12260). The supernatants were collected at 60 h post transfection (hpt), centrifuged at 5000 rpm for 10 min, and filtered. The lentivirus supernatants supplemented with 5 µg/mL polybrene (Sigma-Aldrich) were added into the *ddx21*+/- HeLa cells. The cells were horizontally centrifuged at 1000 rpm for 90 min and incubated at 37°C for 48 h. The supernatants were then replaced with 10% FBS supplemented with blasticidin (Merck-Millipore) for 72 h. The cells were subcloned by limiting dilution and confirmed by immunoblotting. The primer sequences for stable expression are listed in Supplementary Table 1.

### Statistical analysis

Data were expressed as means ± standard deviations. Significance was determined with the two-tailed independent Student’s *t* test (*p*<0.05) between two groups. A one-way ANOVA was followed by Tukey’s to compare multiple groups (>2).

## Results

### DDX21 positively regulates IFN-β signaling pathway

To confirm the role of DDX21 in virus replication as well as its role in anti-viral innate immunity, DDX21 was knocked down followed by VSV infection. The results showed that the knockdown of DDX21 significantly reduced the expression of VSV G protein at 12, 18, and 24 h post infection (hpi) (Fig. 1A). Consistently, the virus titers were significantly impaired after DDX21 knockdown at 6, 12, and 18 hpi (Fig. 1B). To study the role of DDX21 in innate immunity, the expression levels of IFN-β and downstream interferon-stimulate gene (ISG), interferon-induced protein with tetratricopeptide repeats 1 (IFIT1), were evaluated. The results showed that the knockdown of DDX21 inhibited the mRNA levels of IFN-β (Fig. 1C) and IFIT-1 (Fig. 1D). Consistently, phosphorylation of TBK1 was significantly inhibited at 6, 12 and 18 hpi. Similar results were observed in DDX21 knockdown cells after treatment of poly (I:C), confirmed the positive regulation of IFN-β pathway by DDX21 (Fig. 1E-G). For further confirmation, we generated DDX21 knockout cells. Unfortunately, after three rounds of screening, only one (1/225) heterozygous clone (*ddx21+/-*) was identified by sequencing (Fig. S1A). Western blotting showed that compared with WT cells, DDX21 expression was significantly inhibited in ddx21+/- cells (Fig. S1B). The regulation of virus titers and IFN-β pathway was in accord with the knockdown results (Fig. S1C-G). However, using the overexpression model, we observed that transfection of exogenous DDX21 only increased VSV titers at 12 hpi (Fig. 1I), and the synthesis of virus G protein was unchanged at all time points pi (Fig. 1H). Correspondingly, DDX21 overexpression had no effect on IFN-β production or the mRNA levels of ISGs (Fig. 1J–K). Collectively, these results suggested that DDX21 positively regulates IFN-β signaling pathway. However, the discrepancy between the knockdown and overexpression models raised the question of whether, in addition to DDX21 expression, the modification of DDX21 plays a role in anti-viral innate immunity.

**Figure 1.**
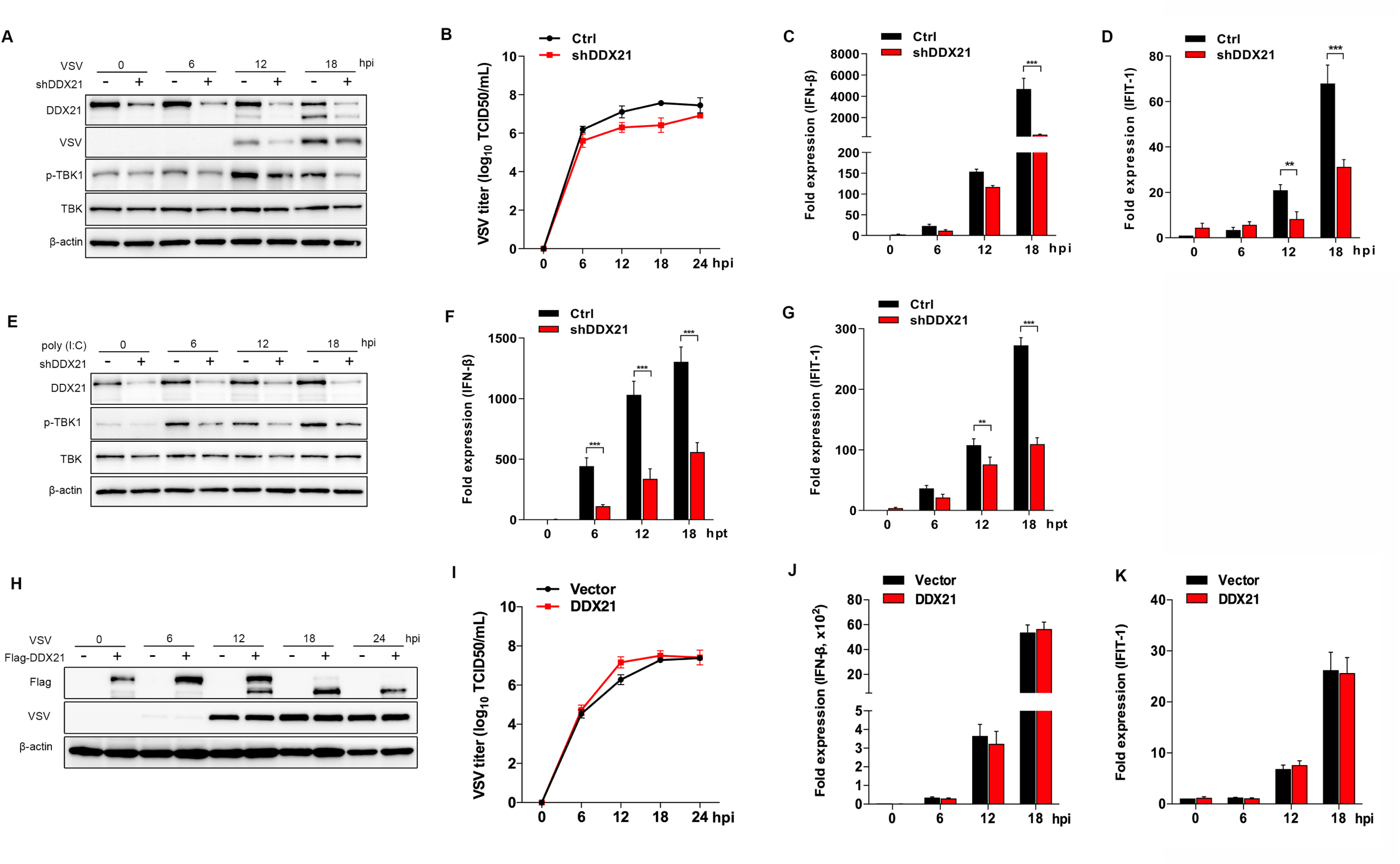
DDX21 positively regulates IFN pathway. (A) HeLa cells with stable knockdown of DDX21 and control cells were mock treated or infected with VSV at an MOI of 1, respectively. Cells were harvested at 6, 12, 18, and 24 hpi, and detected using immunoblot analysis with anti-DDX21, anti-VSV-G, anti-p-TBK1, anti-TBK1 or anti-β-actin antibody. (B) Extracellular virus yields in DDX21 knockdown and control cells. (C-D) Virus infection experiments were performed as in A. Cells were harvested and detected using qRT-PCR with IFN-β (C) and IFIT-1(D) primers, respectively. (E) HeLa cells with stable knockdown of DDX21 and control cells were mock treated or transfected with poly (I:C) (20 μg/mL), respectively. Cells were harvested at 6, 12 and 18 hpt and detected using immunoblot analysis with anti-DDX21, anti-VSV-G, anti-p-TBK1, anti-TBK1 or anti-β-actin antibody. (F-G) Poly (I:C) treatment experiments were performed as in E. Cells were harvested and detected using qRT-PCR with IFN-β (F) and IFIT-1(G) primers, respectively. (H) HeLa cells were transfected with either empty vector or Flag-DDX21. At 24 h after transfection, cells were mock treated or infected with VSV at an MOI of 1. Cells were harvested at 6, 12, 18, and 24 hpi, and detected using immunoblot analysis with anti-DDX21, anti-VSV-G, or anti-β-actin antibody. (I) Extracellular virus yields in empty vector or Flag-DDX21 transfected group. (J-K) Virus infection experiments were performed as in G. Cells were harvested and detected using qRT-PCR with IFN-β (J) and IFIT-1(K) primers, respectively. Data are presented as means from three independent experiments. *, P<0.05, **, P<0.01, ***, P<0.001.

### The cleavage of DDX21 by virus infection and treatment with RNA/DNA ligands

Interestingly, in both the knockdown and overexpression experiments, we observed the apparent cleavage of DDX21 in the course of VSV infection (Fig. 1A and H, Fig. S1C). To further confirm the cleavage of DDX21 by virus infection, cells were infected by two RNA viruses, VSV and NDV, and one DNA virus, HSV-1, respectively, followed by DDX21 detection. As expected, VSV and NDV apparently cleaved DDX21 at 12, 18, and 24 hpi. The full-length DDX21 almost disappeared at 18 and 24 hpi post VSV and NDV infection (Fig. 2A and B). In comparison, HSV-1 only slightly cleaved DDX21, even at the late stage of infection, and the amount of full-length DDX21 was not significantly decreased (Fig. 2C). Statistically, at 18 and 24 hpi, the ratio of DDX21/cleaved DDX21/β-actin was 33-564 in VSV- and NDV-infected cells, compared with 2.4-2.6 in HSV-1-infected cells (Fig. 2D-F). A549, Huh7, and THP-1 cells were then utilized to test whether virus-triggered DDX21 cleavage was cell type-dependent. The results showed that DDX21 was cleaved in A549, Huh7, or THP-1 cells upon VSV or NDV infection (Fig. S2A-D). Additionally, the cleavage of DDX21 was observed in cells infected with SeV, another RNA virus (Fig. S2E).

**Figure 2.**
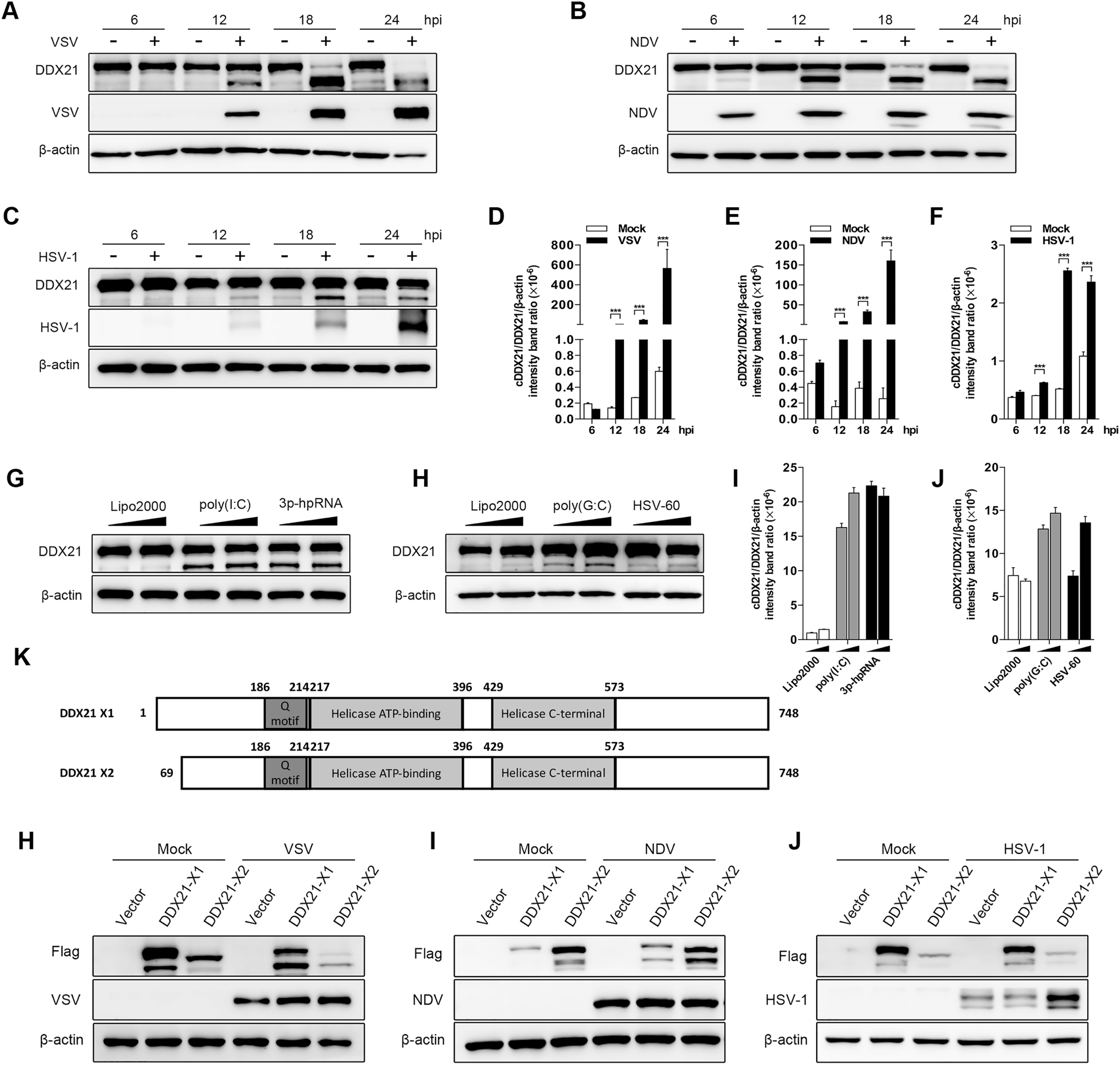
Virus infection or treatment with RNA/DNA ligands leads to the cleavage of DDX21. (A-C) HeLa cells were mock treated or infected with VSV (A), NDV (B) or HSV-1 (C), respectively, at an MOI of 1. Cells were harvested at 6, 12, 18, and 24 hpi, and detected using immunoblot analysis with anti-DDX21, anti-β-actin or anti-viral-protein (VSV-G, NDV-NP, HSV-1-gD) antibody. (D-F) Representative results are shown with graphs representing the intensity band ratio of cleaved DDX21 (cDDX21)/DDX21/β-actin normalized to the control condition for VSV (D), NDV (E) and HSV-1 (F) infection group. Data are presented as means from three independent experiments. ***, P<0.001.

(G-H) HeLa cells were transfected with RNA ligands (poly (I:C) or 3p-hpRNA, G) or DNA ligands (poly (G:C) or HSV-60, H), respectively. At 18 hours post tranfection (hpt), cells were harvested and detected using immunoblot analysis with anti-DDX21 or anti-β-actin antibody.

(I-J) Representative results are shown with graphs representing the intensity band ratio of cleaved DDX21 (cDDX21)/DDX21/β-actin normalized to the control condition for RNA (I) and DNA (J) ligand treatments group.

(K) Schematic representation of two transcript isoforms of DDX21.

(H-J) HeLa cells were transfected with either empty vector, Flag-DDX21-X1, Flag-DDX21-X2. At 24 h after transfection, cells were mock treated or infected with VSV (H), NDV (I) or HSV-1 (J), respectively, at an MOI of 1. Cells were harvested at 18 hpi, and detected using immunoblot analysis with anti-DDX21, anti-β-actin or anti-viral-protein (VSV-G, NDV-NP, HSV-1-gD) antibody.

Given that DDX21 was cleaved upon RNA and DNA virus infection, and as DDX21 belongs to the RNA helicase family that can bind various RNAs (22), we propose that this cleavage is triggered by virus nucleotides. Two RNA ligands, poly (I:C) and 3p-hpRNA, and two DNA ligands, poly (G:C) and HSV-60, were used to evaluate their role in DDX21 cleavage. As expected, both RNA and DNA ligands cleaved DDX21 (Fig. 2G and H), and higher cleavage efficiency was observed upon treatment with RNA ligands, compared with that of DNA ligands (Fig. 2I and J), which is in accordance with the results of virus infection. There are two transcript variants of DDX21, isoform 1 (X1), which encodes the full-length of DDX21, and isoform 2 (X2), with a shorter N-terminus (Δ1-86) compared to X1. Here, we showed that both exogenously expressed DDX21 X1 and X2 were cleaved upon VSV, NDV, and HSV-1 infection (Fig. 2H-J). Collectively, these results clearly demonstrated that DDX21 was cleaved by virus infection and treatment with RNA/DNA ligands.

### DDX21 was cleaved in a caspase-dependent manner

To further confirm whether DDX21 was cleaved or degraded upon virus infection and RNA/DNA ligand treatment, cells were treated with caspase inhibitor z-VAD-FMK, neddylation inhibitor MLN4924, proteasome inhibitor MG-132, and autophagy inhibitors wortmannin and CQ, respectively, followed by virus or treatment with ligand. The results showed that DDX21 cleavage was completely inhibited after treatment with the caspase inhibitor z-VAD-FMK in cells infected with VSV, NDV, and HSV-1 (Fig. 3A–D, lane 4) or treated with RNA ligand poly (I:C) (Fig. 3D, lane 4). Notably, MG132 and CQ also seem to inhibit the cleavage of DDX21, especially in VSV- and NDV-infected cells (Fig. 3A and B, lanes 6 and 7). However, the expression of viral protein was also inhibited after treatment with these two drugs, indicating that the inhibition of DDX21 cleavage may be due to their inhibition of virus replication. In comparison, z-VAD-FMK had no effect on virus replication. These results indicated that DDX21 was cleaved in a caspase-dependent manner.

**Figure 3.**
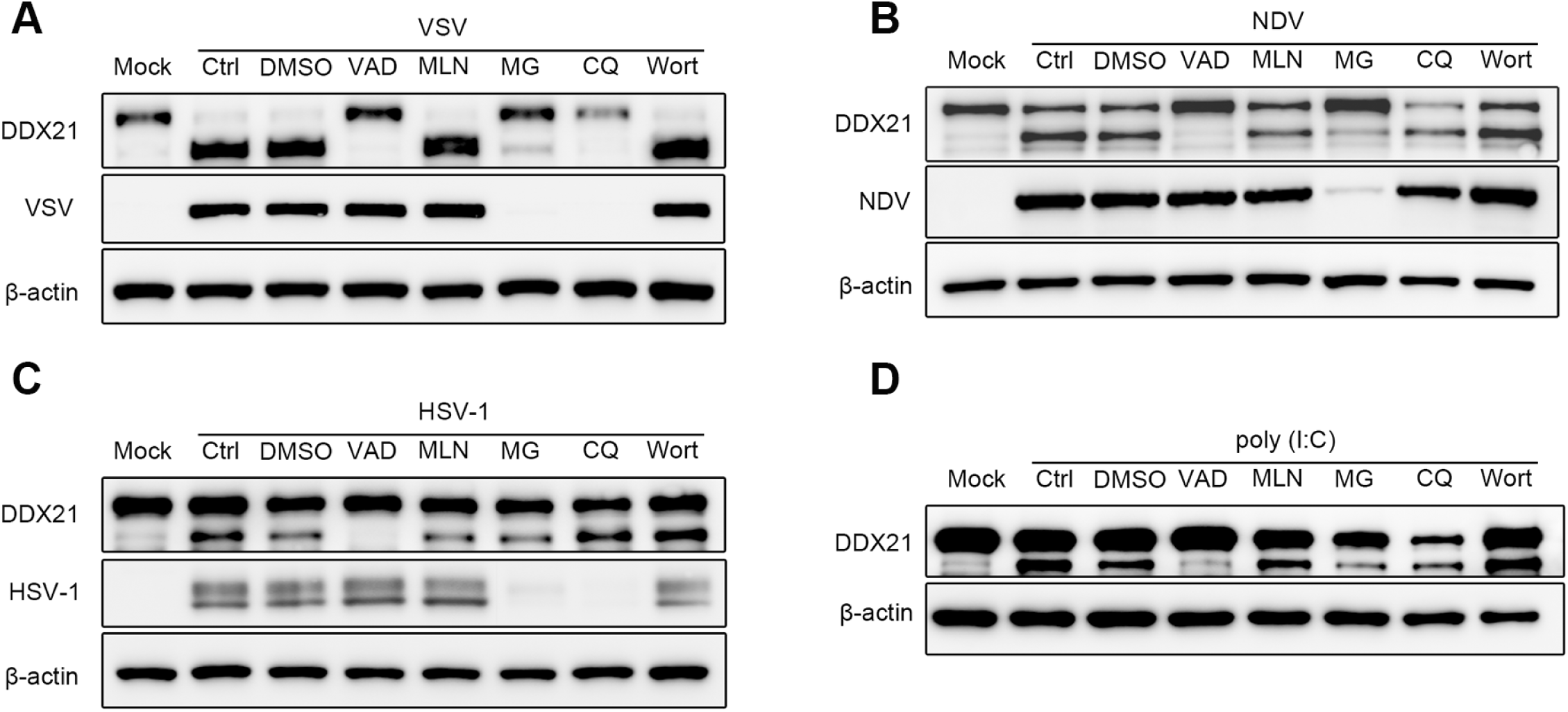
The caspase-dependent cleavage of DDX21 triggered by virus infection or treatmens of RNA ligands. (A-C) HeLa cells were mock infected or infected with VSV(A), NDV (B) or HSV-1 (C) at an MOI of 1 and maintained in the presence of DMSO control, z-VAD-FMK (VAD), MLN4924 (MLN), MG132 (MG), CQ or wortmannin (Wort) for 18 h. Cells were harvested and detected using immunoblot analysis with with anti-DDX21, anti-β-actin or anti-viral-protein (VSV-G, HSV-1-gD, NDV-NP) antibody. (D) HeLa cells were mock treated or transfected with RNA ligands (poly (I:C) and maintained in the presence of DMSO control, z-VAD-FMK, MLN4924, MG132, CQ or wortmannin for 18 h. Cells were harvested and detected using immunoblot analysis with with anti-DDX21 or anti-β-actin antibody.

### Caspase-3/6 cleaved DDX21 at D^126^ in response to VSV infection

DDX21 is characterized by several known domains, Q motif (amino acid (aa) 186-214), helicase ATP-binding domain (aa 217-396), and helicase C-terminal domain (aa 429-573) (Fig. 4A). To characterize the cleavage sites of DDX21, based on the identified domains, several deletion mutants (Δ1-216, Δ217-395, Δ396-573, Δ574-784) were generated to test the critical domain involved in DDX21 cleavage. As shown in Fig. 4B, deletion of 1-216 abrogated the cleavage of DDX21. Notably, unlike endogenous DDX21 expression, exogenous expression of DDX21 truncates alone is able to induce cleavage to some extent (Fig. 4B, left panel). Similar results could be observed in the overexpression of wild-type (WT) DDX21 (Fig. 2H-J). Combined with the results showing that aa 1-216 is critical for DDX21 (Fig. 4B) and aa 1-86 is not required for DDX21 cleavage (Fig. 2H-J), the cleavage sites are within aa 87-216 of DDX21. The caspase cleavage sites were then predicted by CaspDB (http://caspdb.sanfordburnham.org). The results showed that all three aspartates (Asp, D), D87, D126, D160, are the putative caspase cleavage sites (Fig. 4C). Therefore, DDX21 with single, double, and triple mutations of these three Asp sites were generated, respectively, followed by transfection and virus infection. The results showed that D126A, but not D87A or D160A, is sufficient and necessary for DDX21 cleavage (Fig. 4D, lanes 4, 6, 8 and 9). The truncates of DDX21 1-125 and 127-784, together with WT DDX21, were transfected into cells to further confirm that the cleavage was mediated by D126. As expected, WT, but not 1-125 and 127-784 DDX21, was cleaved upon virus infection (Fig. 4E). It should be noted that based on the molecular weight of DDX21, we proposed that the “cDDX21” we observed in endogenous cleavage (∼73kD) was DDX21 127-784, while the relatively small cleaved DDX21 (∼14kD) was degraded upon virus infection (Fig. 4E). Previous reports showed that the cleavage sites specific for caspases had a general motif (37). The motif for DDX21 cleavage is Glu-Ile-Asp (E-I-D), which is the putative substrate for caspase-3 or caspase-6 (Fig. 4F). *casp3-/-* and *casp6*-/- cells were generated to verify their role in DDX21 cleavage. As expected, knockout of *casp3* and especially *casp6* significantly inhibited the cleavage of DDX21 (Fig. 4G-H). DDX21 cleavage was almost blocked in *casp3/6* double knockout cell lines (Fig. 4I). These results indicated that virus infection cleaved DDX21 at D126 via caspase-3/6.

**Figure 4.**
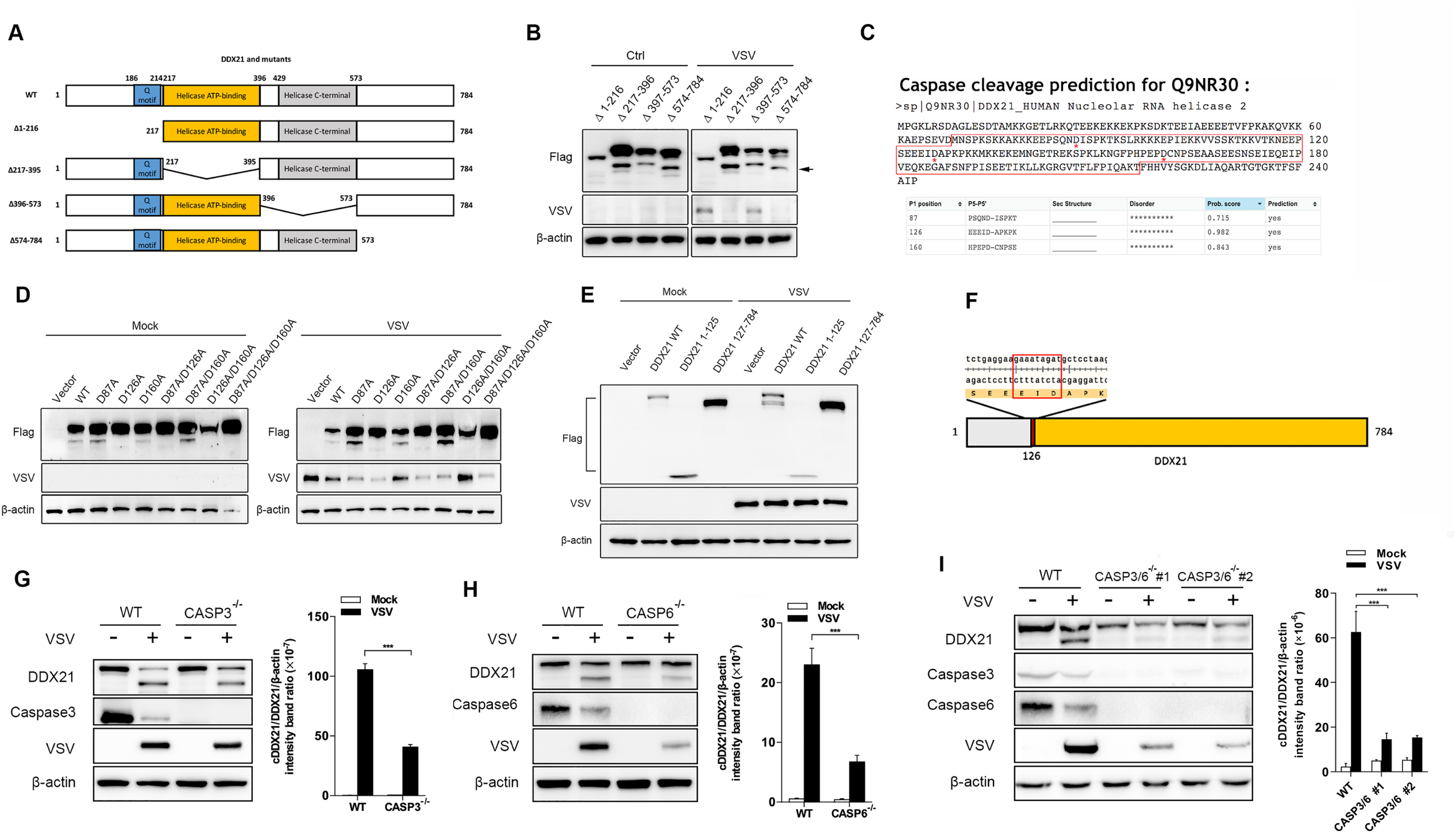
DDX21 was cleaved at D^126^ by caspase-3/6 in response to VSV infection. (A) Schematic representation of WT DDX21 and its deletion mutants. (B) HeLa cells were transfected with Flag tagged DDX21 truncates (Δ1-216, Δ217-395, Δ396-573, Δ574-784). At 24 h after transfection, cells were mock treated or infected with VSV at an MOI of 1. Cells were harvested at 18 hpi, and detected using immunoblot analysis with anti-Flag, anti-VSV-G, or anti-β-actin antibody. (C) The prediction results of caspase cleavage sites for DDX21 based on CaspDB (http://caspdb.sanfordburnham.org). The red frame indicates aa 87-216 of DDX21. The red asterisks (*) indicate the putative caspase cleavage sites. (D) HeLa cells were transfected with either empty vector or Flag tagged WT and mutants of DDX21 (D87A, D126A, 160A, D87A/D126A, D87A/160A, D126A/D160A, D81A/D126A/160A). At 24 h after transfection, cells were mock treated or infected with VSV at an MOI of 1. Cells were harvested at 18 hpi, and detected using immunoblot analysis with anti-Flag, anti-VSV-G, or anti-β-actin antibody. (E) HeLa cells were transfected with either empty vector or Flag tagged WT and truncates of DDX21 (1-125, 127-784). At 24 h after transfection, cells were mock treated or infected with VSV at an MOI of 1. Cells were harvested at 18 hpi, and detected using immunoblot analysis with anti-DDX21, anti-VSV-G, or anti-β-actin antibody. (F) Schematic representation of amino acids around cleavage site. The red frame indicates the general motif for caspase cleavage. (G-I) WT and *casp3* (G), *casp6* (H) knockout and casp3/6 double-knockout (I) HeLa cells were mock treated or infected with VSV at an MOI of 1, respectively. Cells were harvested at 18 hpi, and detected using immunoblot analysis with anti-DDX21, anti-caspase-3/6, anti-VSV-G, or anti-β-actin antibody. Representative results are shown with graphs representing the intensity band ratio of cleaved DDX21 (cDDX21)/DDX21/β-actin normalized to the control condition.

### DDX21 was cleaved and translocated from the nucleus to the cytoplasm in response to virus infection

DDX21 was identified to be a nucleolar helicase that is required for pre-rRNA processing (22, 23, 38). Other reports showed that DDX21 is localized with DDX1, DDX26, and TRIF to sense dsRNA in the cytosol (20). To study whether DDX21 localization was affected by virus infection, cells were infected with VSV and HSV-1, followed by IFA test. As expected, endogenous DDX21 was predominantly localized in the nucleolus in mock-infected cells (Fig. 5A and C, upper panels). DDX21 was translocated from the nucleolus to nucleoplasm and cytoplasm after VSV and HSV-1 infection (Fig. 5A and C, middle and lower panel). Statistically, at 12 and 24 hpi, cells with cytoplasmic DDX21 accounted for 23% and 68% of all cells for VSV (Fig. 5B) and 24% and 49% of all cells for HSV-1 (Fig. 5D), respectively, indicating that VSV could induce DDX21 translocation more efficiently than HSV-1. Next, we evaluated whether the cleavage of DDX21 affected its localization. WT, D126A, 1-125, and 127-784 DDX21 (Fig. 5E), together with empty vector, were transfected into cells followed by VSV infection. The results showed that in mock-infected cells, WT, D126A, and 127-784 DDX21 predominantly localized in the nucleolus (Fig. 5F, left panel). WT and 127-784, but not D126A DDX21, efficiently translocated from the nucleolus to nucleoplasm and cytoplasm after VSV infection (Fig. 5F and G), indicating that blockage of DDX21 cleavage inhibited its translocation. Moreover, the statistical analysis results showed the most efficient translocation of DDX21 cleaved form (cDDX21, 126-784), further indicating DDX21 cleavage promotes its translocation from nucleus to cytoplasm. Interestingly, 1-125 DDX21 was diffusely localized in the nucleus and cytoplasm in both mock- and VSV-infected cells (Fig. 5F and G). Collectively, these results demonstrated that the virus induced the translocation of DDX21, which is affected by DDX21 cleavage.

**Figure 5.**
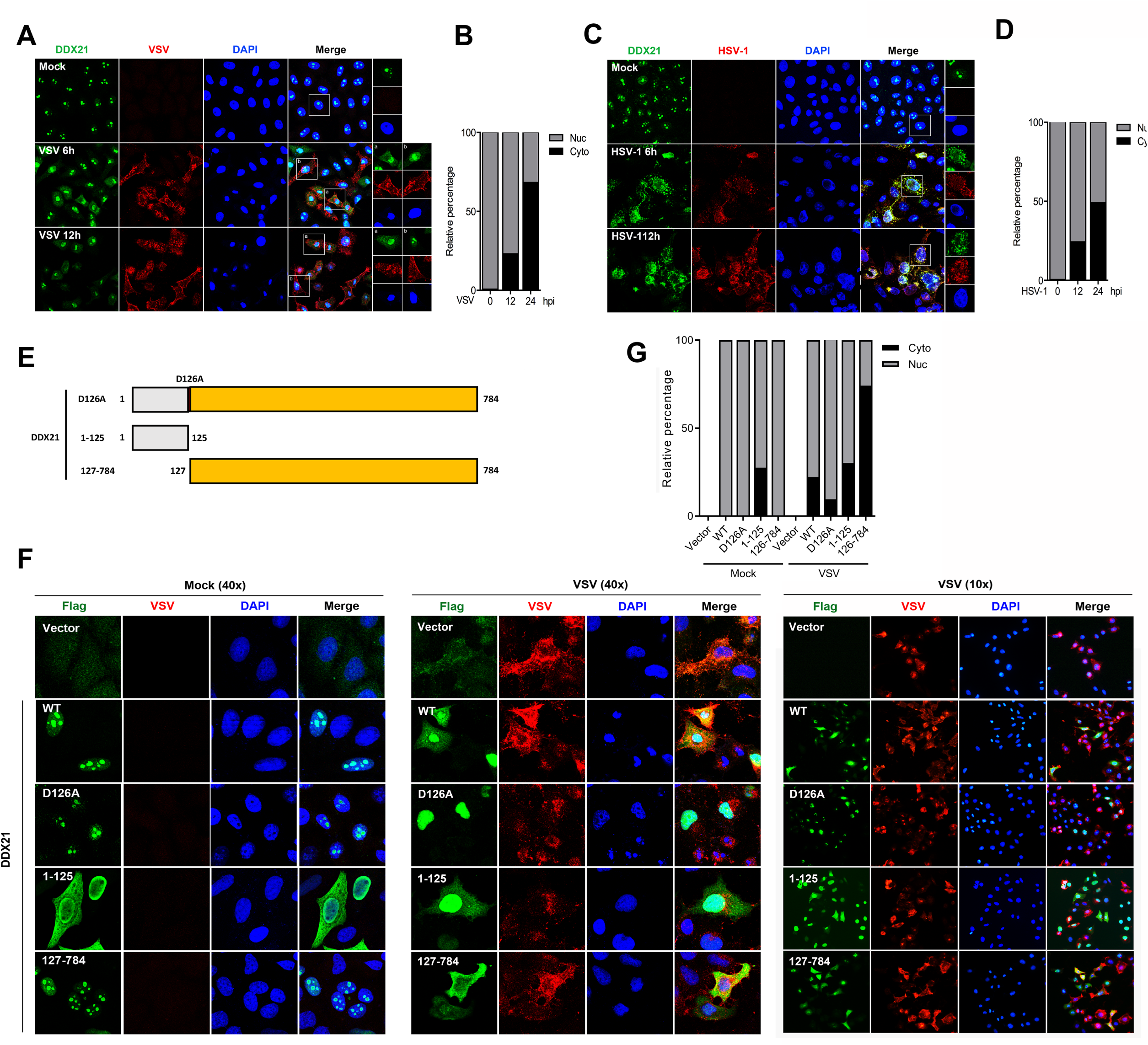
Virus infection triggers the translocation of DDX21 from nucleus to cytoplasm which is dependent on its cleavage. (A, C) HeLa cells were mock treated or infected with VSV (A) or HSV-1 (C) at an MOI of 1. At 6 and 12 hpi, cells were fixed, and processed for IF using anti-DDX21 or anti-viral-protein (VSV-G, HSV-1-gD) antibody. Nuclei were stained with 1 μg/mL of DAPI. (B, D) The relative percentages of cells with nuclear and cytoplasmic DDX21 staining upon VSV (B) or HSV-1 (D) infection were quantified. 10 images in 20 high-powered fields (HPFs) were obtained randomly in different fields. (F) HeLa cells were transfected with either empty vector or Flag tagged WT, D126A, 1-126 or 127-784 DDX21. At 24 h after transfection, cells were mock treated or infected with VSV at an MOI of 1. At 12 hpi, cells were fixed, and processed for IF using anti-Flag or anti-VSV-G antibody. Nuclei were stained with 1 μg/mL of DAPI. (G) The relative percentages of cells with nuclear and cytoplasmic DDX21 staining in cells transfected with either empty vector or Flag tagged WT, D126A, 1-126 or 127-784 DDX21 upon virus infection were quantified. 10 images in 20 high-powered fields (HPFs) were obtained randomly in different fields.

### DDX21 cleavage led to the inhibition of the IFN-β signaling pathway

Given that the blockage of DDX21 inhibits its translocation, we next aimed to explore the effect of DDX21 cleavage in the regulation of innate immunity. *Ddx21+/-* cells were transfected with WT and truncated DDX21, followed by the detection of virus replication and IFN-β signaling pathway. Interestingly, transfection of WT and truncated DDX21 did not affect virus replication, evidenced by viral protein expression and titers in the supernatants (Fig. 6A and B). In contrast, although WT DDX21 has no effect, the intact form of DDX21 (D126A) did increase the mRNA levels of IFN-β and ISGs (IFIT-1 and MX1) after VSV infection. More importantly, transfection of cDDX21 significantly inhibited IFN-β signaling pathway (Fig. 6C-E). These results were further confirmed using poly (I:C) as the stimulator (Fig. 6F-H). To identify whether DDX21 cleavage regulates IFN-β signaling pathway in a time-dependent manner, *ddx21+/-* cells stably expressing flag-tagged WT and D126A DDX21 infected by VSV or treated with poly (I:C) and collected at different time points. DDX21 was cleaved in WT but not D126A DDX21-expressed cells. However, no difference was observed in viral protein synthesis between WT and D126A DDX21-expressed cells (Fig. 6I). More importantly, blockage the cleavage (D126A) significantly increased the IFN-β promoter activity (Fig. 6K and O), mRNA levels of IFN-β and ISGs (IFIT-1 and MX1) (Fig. 6L–N, P-R) and IFN-β production in the supernatants (Fig. 6S) especially at the late stage of virus infection and poly (I:C) treatment. These results clearly indicated that DDX21 cleavage inhibits the IFN-β signaling pathway.

**Figure 6.**
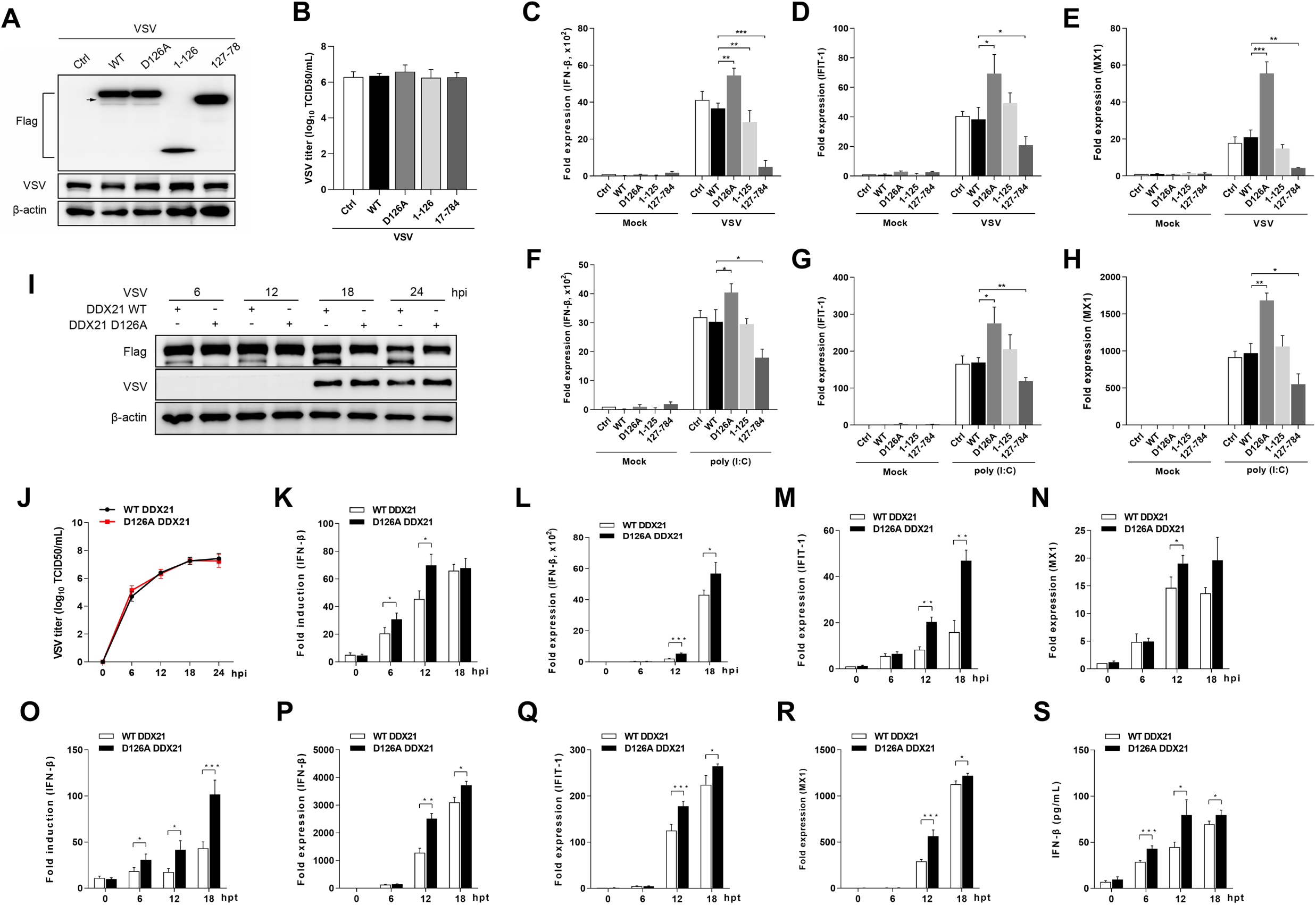
Cleavage of DDX21 impairs IFN-β production. (A) *ddx21+/-* HeLa cells were transfected with either empty vector or Flag tagged WT, D126A, 1-126 or 127-784 DDX21. At 24 h after transfection, cells were mock treated or infected with VSV at an MOI of 1. At 18 hpi, cells were harvested and detected using immunoblot analysis with anti-Flag, anti-VSV or anti-β-actin antibody. The arrow indicates cleaved DDX21. (B) Extracellular virus yields in WT and truncated DDX21 transfected *ddx21+/-* HeLa cells. (C-E) Virus infection experiments were performed as in A. Cells were harvested and detected using qRT-PCR with IFN-β (C), IFIT-1(D) and MX1 (E) primers, respectively. (F-H) *ddx21+/-* HeLa cells were transfected with either empty vector or Flag tagged WT, D126A, 1-126 or 127-784 DDX21. At 24 h after transfection, cells were mock treated or transfected with poly (I:C) (20 μg/mL), respectively. Cells were harvested at 12 hpt and detected using qRT-PCR with IFN-β (F), IFIT-1 (G), or MX1 (H) primers. (I) *ddx21+/-* cells stably expressing flag-tagged WT and D126A DDX21 were mock treated or infected with VSV at an MOI of 1, respectively. Cells were harvested at 6, 12, 18 and 24 hpi, and detected using immunoblot analysis with anti-DDX21, anti-VSV-G, or anti-β-actin antibody. (J) Extracellular virus yields in *ddx21+/-* cells stably expressing flag-tagged WT and D126A DDX21 (K) *ddx21+/-* cells stably expressing flag-tagged WT and D126A DDX21 were co-transfected with IFN-β-Luc and PRL-TK. At 12 hpt, cells were mock treated or infected with VSV at an MOI of 1, respectively. Cells were harvested at 6, 12 and 18 hpi and assessed for luciferase activity. The results are presented as relative luciferase activity. (L-N) *ddx21+/-* cells stably expressing flag-tagged WT and D126A DDX21 were mock treated or infected with VSV at an MOI of 1, respectively. Cells were harvested at 6, 12 and 18 hpi and detected using qRT-PCR with IFN-β (L), IFIT-1 (M), or MX1 (N) primers. The results are presented as relative luciferase activity. (O) *ddx21+/-* cells stably expressing flag-tagged WT and D126A DDX21 were co-transfected with p-125Luc and PRL-TK. At 12 hpt, cells were mock treated or transfected with poly (I:C) (20 μg/mL). Cells were harvested at 6, 12 and 18 hpt and assessed for luciferase activity. The results are presented as relative luciferase activity. (P-R) *ddx21+/-* cells stably expressing flag-tagged WT and D126A DDX21 were mock treated or transfected with poly (I:C) (20 μg/mL), respectively. Cells were harvested at 6, 12 and 18 hpt and detected using qRT-PCR with IFN-β (P), IFIT-1 (Q), or MX1 (R) primers. (S) Poly (I:C) treatments were performed as in O. Supernatants were collected for quantization of IFN-β by ELISA. Data are presented as means from three independent experiments. *, P<0.05, **, P<0.01, ***, P<0.001.

### Cleavage of DDX21 inhibited the formation of DDX1-DDX21-DHX36 complex

Given that DDX21 was reported to act as the scaffold protein in the complex of three dsRNA-sensing helicases (DDX1-DDX21-DHX36) (20, 39) (Fig. 7D), we hypothesized that DDX21 cleavage attenuates the inhibition of the IFN-β signaling pathway by inhibition of the formation of DDX1-DDX21-DHX36 complex. Since DDX21 was reported to exist as a homodimer (25, 27), the cleavage of DDX21 on its self-interaction was studied first. As shown in Fig. 7A, using full-length DDX21 as the bait, the full-length and 127-784, but not 1-125 DDX21, could be immunoprecipitated (Fig. 7A). In accordance, 127-784 DDX21 was interacted with 127-784 and full-length, but not 1-125 DDX21 (Fig. 7B). In contrast, 1-125 DDX21 was not immunoprecipitated with WT and truncated DDX21 (Fig. 7C). These results indicated that DDX21 interacted with itself through its C-terminal 127-784. Next, we evaluated whether DDX21cleavage inhibits the formation of DDX1-DDX21-DHX36 complex. The results showed that compared with intact DDX21 (D126A), cDDX21 (127–784) has a lower affinity for both DDX1 and TRIF (Fig. 7E-F). In comparison, D126A, 127-784 and 1-125 DDX21 showed similar binding ability to DHX36 (Fig. 7G). Collectively, these results indicated cleavage of DDX21 inhibited the formation of DDX1-DDX21-DHX36 complex and its interaction to downstream adaptor TRIF.

**Figure 7.**
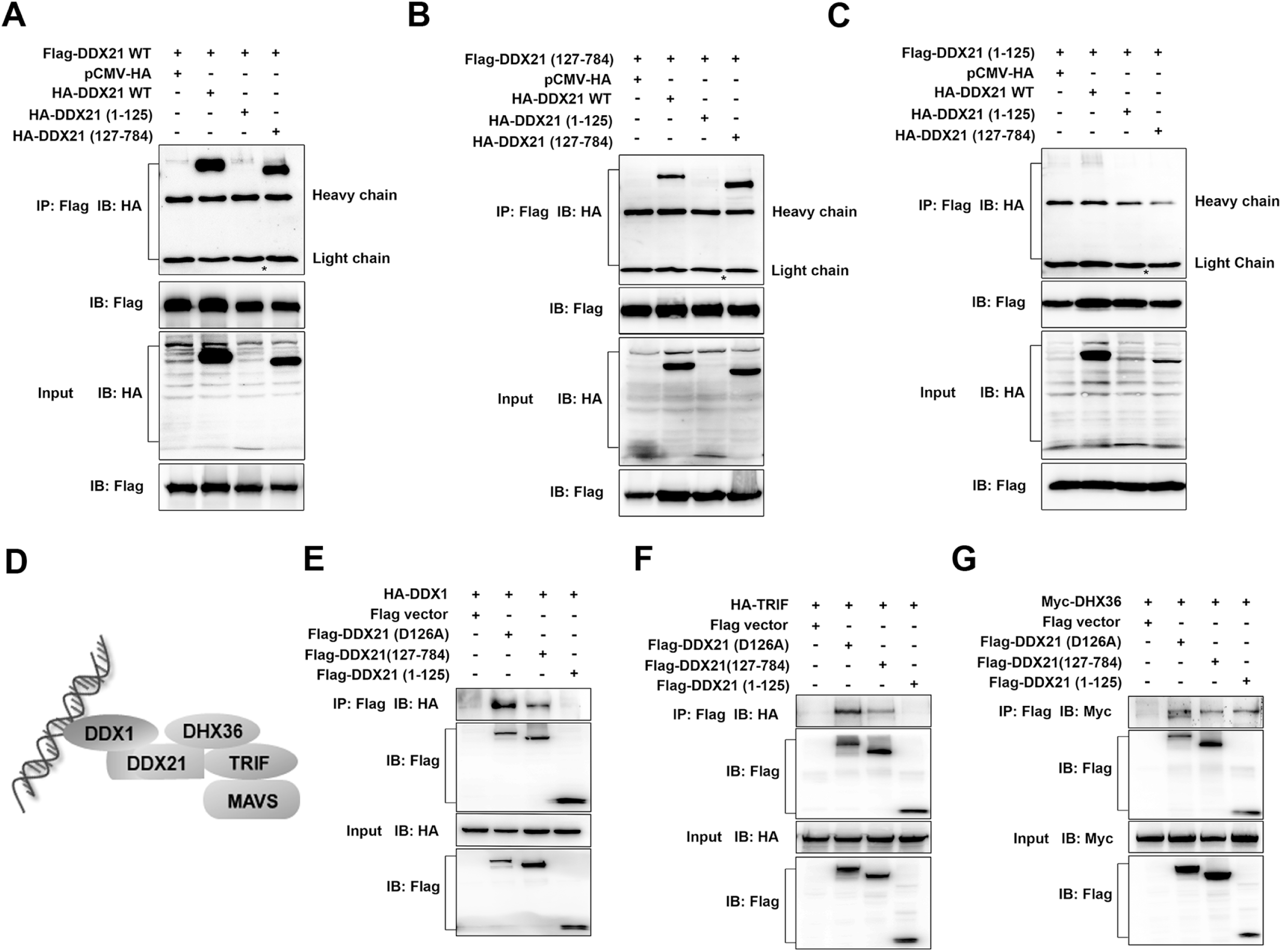
DDX21 cleavage inhibits its interaction with DDX1 and TRIF. (A-C) HeLa cells were transfected with Flag tagged WT (A), 127-784 (B), 1-125 (C) DDX21, respectively, together with HA tagged WT, 1-125 or 127-784 DDX21. At 24 hpt, cells were harvested, immunoprecipitated with anti-Flag antibody, and further detected using immunoblot analysis with anti-HA or anti-Flag antibody. Expression levels of the proteins were analyzed by immunoblot analysis of the lysates with anti-HA or anti-Flag antibody. The asterisk (*) indicates the predicted size of HA-DDX21 (1-125). (D) The existing model of foreign RNA recognition by DDX1-DDX21-DDX36 complex. (E-G) HeLa cells were co-transfected with Flag-tagged D126A, 1-125, 127-784 DDX21 or empty vector, and HA-tagged DDX1 (E), HA-tagged TRIF (F) and Myc-tagged DHX36 (G), respectively. At 24 hpt, cells were harvested, immunoprecipitated with anti-Flag antibody, and further detected using immunoblot analysis with anti-HA or anti-Flag antibody. Expression levels of the proteins were analyzed by immunoblot analysis of the lysates with anti-HA or anti-Flag antibody.

## Discussion

Intensive functional and structural research over the years has clearly demonstrated that RLRs selectively bind viral RNA ligands and trigger downstream signaling (40). Several other DExD/H-box helicases have been implicated in anti-viral innate immunity, but the fundamental questions such as the precise regulation mechanism how DExD/H-box helicases regulate innate immunity remain to be elucidated. Here, we demonstrated that DDX21 is cleaved at D^126^ after virus infection and RNA/DNA ligand treatment via the caspase-3/6 pathway, leading to the inhibition of immune responses.

To date, there has been limited research on the cleavage of RNA helicases. Human RNA helicase A, a nuclear helicase that unwinds dsRNA, dsDNA, and RNA:DNA duplexes, was reported to be cleaved by caspase-3 during apoptosis (41). Another study also showed that the CARD-containing helicase (Helicard) is cleaved by caspases upon apoptotic stimuli (42). However, no studies have shown that the cleavage of DNA/RNA helicases is involved in innate immunity and virus infection. Our data showed the obvious cleavage of DDX21 in response to virus infection and treatment with ligands. The fact that three RNA viruses and one DNA virus, together with two RNA and two DNA ligands, trigger varying degrees of DDX21 cleavage leads us to speculate that the ability of DDX21 cleavage may be shared by most viruses.

The caspase family of cysteine proteases is involved in apoptosis and innate immune signaling (43). In terms of the innate immunity pathway, caspase-1 is one of the most well-studied caspases, which is able to cleave proIL-1b and proIL-18 and triggers inflammasomes (44). It has also been reported that caspases are involved in the RLR-mediated type I IFN response. For instance, caspase-12 positively modulates the IFN-β signaling pathway by regulating E3 ubiquitin ligase TRIM25-mediated ubiquitination of RIG-I (45). Activated caspase-3/7/9 suppress mitochondrial DNA-induced stimulator of interferon genes protein (STING)-mediated type I IFN production (46). Here, we showed that the cleavage of DDX21 was completely recovered in the presence of caspase inhibitor z-VAD-FMK (Fig. 3), and the cleavage was probably mediated by caspase-3/6 (Fig. 4). It should be noted that knockout of *casp3* and *casp6* did not completely blocked DDX21 cleavage, indicating that the involvement of other caspases could not be completely excluded. A recent study revealed that caspase-6 cleaves IL-1R-associated kinase-M (IRAK-M) and reduces IκBα degradation, thereby increasing TNF-α production (47). The function of caspase-6 in the type I IFN pathway has not been reported. Our study indicated that caspase-3/6 cleaves DDX21 and thus likely regulates the IFN-β signaling pathway.

In a resting state, DDX21, together with its binding partners, c-Jun, WDR46, SIRT7, etc., is localized in the cell nucleolus (23, 38, 48). The nucleolar localization of DDX21 is necessary for its pre-rRNA processing and RNA unwinding (23, 48). Studies have also suggested that DDX21 is involved in innate immunity in the cytoplasm. For example, the DDX1-DDX21-DHX36-TRIF complex may translocate to the mitochondria upon poly(I:C) stimulation (20). The infection of A549 cells with the dengue virus causes DDX21 to partially relocate from the nucleus to the cytoplasm (29). Here, we confirmed that DDX21 localized in the cell nucleolus in mock-infected cells. Virus infection effectively triggered DDX21 translocation from the nucleolus to the cytoplasm. In resting cells, WT, D126A, and 127-784 DDX21 showed nucleolar localization. It is interesting that 1-125 DDX21 diffusely localized in the nucleus and the cytoplasm. Previous reports showed that the 731–740 at the C-terminus of DDX21 interacted with c-Jun, and depletion of c-Jun promotes DDX21 translocation from the nucleolus to the nucleoplasm (23). Therefore, it is possible that the deletion of the C-terminus of DDX21 abolishes the interaction with its binding partner and thus alters its nucleolar localization. Our results that (1) blockage of DDX21 cleavage inhibits DDX21 translocation, (2) the most efficient translocation of cDDX21 upon virus infection, indicated that cleavage of DDX21 promotes its translocation from nucleus to cytoplasm.

Various DDX and DHX helicases play important roles in maintaining the stability of the cell genome (22, 48, 49). To date, the knockout of *ddx21* has never been reported in cells or mice. Studies on other DDXs have also shown that certain DDXs are critical for mouse growth, and knockout of the *ddx* gene results in early embryonic lethality (50, 51). Therefore, DDX21 may also be critical for cell and mouse survival. Nevertheless, DDX21 expression was significantly inhibited in *ddx21+/-* cells. Previous studies and our results demonstrated that depletion of DDX21 significantly inhibited type I IFN production (20), suggesting that DDX21 positively regulate innate immunity. Interestingly, depletion of DDX21 also impaired virus replication. Considering DDX21 is a multifunctional protein which also plays a important role in maintaining the stability of the cell genome (27, 48, 52), it may also influence virus replication in other ways except for innate immunity. Nevertheless, it is undoubtful that DDX21 *per se* plays a positive role in innate immunity. Most importantly, cleavage of DDX21 inhibits innate immunity, but not affects virus infection, which lead us to speculate the cleavage of DDX21 was driven by host for a late counter-regulatory effect to temper immune responses.

Although type I IFN is widely reported to play an essential role against viral infection, the aberrant production of cytokines leads to unexpected pathological consequences in a variety of autoimmune diseases (53, 54). Therefore, the balance between these key pathways is essential for immune homeostasis. Indeed, DNA sensor cGAS has been reported as a key driver of lethal autoimmune disease in the Trex1-deficient mouse model of Aicardi–Goutieres Syndrome (AGS) (55). The importance of excess RLR-dependent signaling, which leads to an IFN signature in the pathogenesis of many autoimmune diseases, such as AGS and systemic lupus erythematosus, has also been clarified (56). These reports highlight the importance of fine-tuning the regulation of the type I IFN signaling pathway. Here, we provided several lines of evidence to demonstrate that host promotes DDX21 cleavage to temper immune responses. (i) the cleavage of DDX21 was observed at the late stage of infection; (ii) the cleavage was a universal phenomenon not only for virus but also for RNA and DNA mimics. (iii) blockage of DDX21 (D126A) cleavages increase IFN production and cleaved DDX21 (127-784aa) inhibits IFN production. (iv) cleavage of DDX21 did not affect virus replication at the late stage of infection. (v) cleavage of DDX21 inhibited the formation of DDX1-DDX21-DHX36 complex. Collectively, from our original and additional data, we inferred that host promotes DDX21 cleavage via caspase3/6 to suppress DDX1-DDX21-DHX36 complex formation for a late counter-regulatory effect to temper immune responses (Fig. 8). An improved understanding of these processes could shed light on the causes of infectious disease and, plausibly, immune disorders involving excessive inflammatory immune activities.

**Figure 8.**
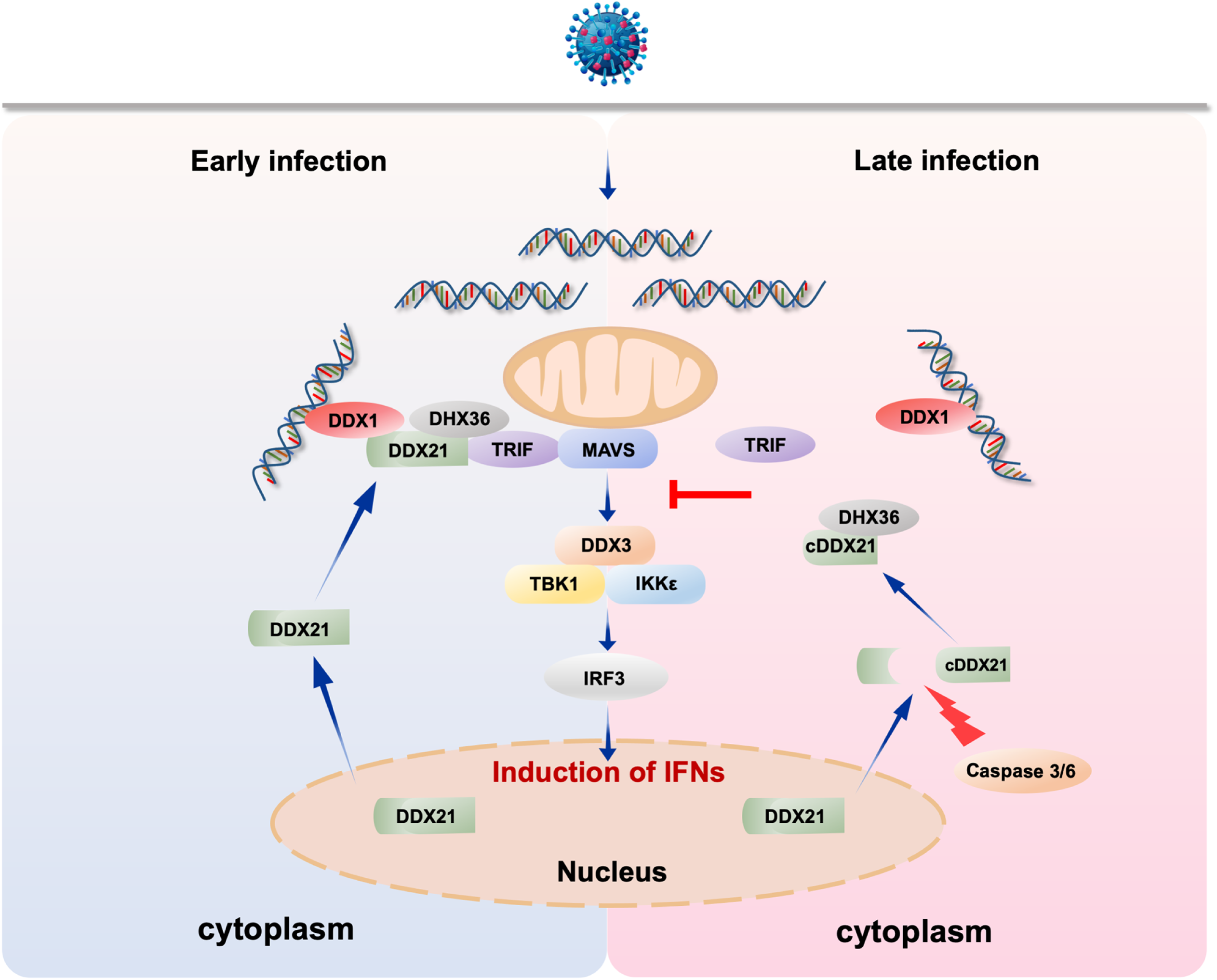
Proposed model for the regulation of innate immunity by DDX21 cleavage during virus infection. The double-edged sword role of DDX21 in regulation of innate immunity upon virus infection. At the early stage of infection, foreign RNA was recognized by DDX1 and recruit DDX21 and DHX36 to form a complex to mediate downstream anti-viral innate immunity. At the late stage of infection, to avoid excessive immune response, host promotes DDX21 cleavage via caspase3/6. Cleaved DDX21 tends to translocate from nucleus to cytoplasm. DDX21 cleavage reduces the interaction between upstream DDX1 and downstream TRIF, and therefore suppresses the signal transduction for a late counter-regulatory effect to temper immune responses.

## Acknowledgements

We thank Dr. Takashi Fujita (Kyoto University) for providing plasmid IFN-β-Luc. We thank Dr. Yasushi Kawaguchi (university of Tokyo, Japan) for providing HSV-1. We thank Dr. Jianchao Wei (Shanghai Veterinary Research Institute, China) for providing VSV. We thank Quan Zhang (Yangzhou University, China) for providing Sendai virus (SeV).

## Supporting Information Legends

**S1 Table. Primers and siRNAs used in this study**

**S1 Figure. DDX21 knockout inhibits IFN signaling pathway**

(A) Confirmation of the genome editing by sequencing of the PCR amplicon from the DDX21 genome of the cell lines.

(B) WT and *ddx21+/-* HeLa cells were seeded in 6-well plates and collected for immunoblot analysis with anti-DDX21, or anti-β-actin antibody.

(C) WT and *ddx21+/-* HeLa cells were mock treated or infected with VSV at an MOI of 1, respectively. Cells were harvested at 6, 12 and 18 hpi, and detected using immunoblot analysis with anti-DDX21, anti-VSV-G, or anti-β-actin antibody.

(D) Extracellular virus yields in DDX21 knockdown and control cells.

(E-G) Virus infection experiments were performed as in C. Cells were harvested and detected using qRT-PCR with IFN-β (E), IFIT-1(F) and MX1 (G) primers, respectively.

**S2 Figure. The cleavage of DDX21 by virus infection in various cell types**

(A-B) A549 cells were mock treated or infected with VSV (A) or NDV (B), respectively, at an MOI of 1. Cells were harvested at 6, 12, 18, and 24 hpi, and detected using immunoblot analysis with anti-DDX21, anti-β-actin or anti-viral-protein (VSV-G, NDV-NP) antibody.

(C-D) Huh7 (C) or THP-1 (D) cells were infected with VSV. The virus infection experiments were performed as in A.

(E) HeLa cells were infected with Sendai virus (SeV). The virus infection experiments were performed as in A.

**S3 Figure. Subcellular distribution of WT and truncated DDX21 by nucleocytoplasmic separation assay**

*ddx21+/-* HeLa cells were transfected with either empty vector or Flag tagged WT, D126A, 1-126 or 127-784 DDX21. At 24 h after transfection, cells were mock treated or infected with VSV at an MOI of 1. At 18 hpi, cells were harvested for nucleocytoplasmic separation assay, and detected using immunoblot analysis with anti-Flag, anti-β-actin or anti-LaminB1 antibody.

